# Time-varying Functional Connectivity Predicts Fluctuations in Sustained Attention in a Serial Tapping Task

**DOI:** 10.1101/2023.06.22.546092

**Authors:** Dolly T. Seeburger, Nan Xu, Marcus Ma, Sam Larson, Christine Godwin, Shella D. Keilholz, Eric H. Schumacher

## Abstract

The mechanisms for how large-scale brain networks contribute to sustained attention is unknown. Attention fluctuates from moment to moment and this continuous change is consistent with dynamic changes in functional connectivity between brain networks involved in the internal and external allocation of attention. In this study, we investigated how brain network activity varied across different levels of attentional focus (i.e., “zones”). Participants performed a finger-tapping task, and guided by previous research, in-the-zone performance or state was identified by low reaction time variability and out-of-the-zone as the inverse. Employing a novel method of time-varying functional connectivity, called the quasi-periodic pattern analysis (i.e., reliable network-level low-frequency fluctuations), we found that the activity between the default mode network (DMN) and task positive network is significantly more anti-correlated during in-the-zone states versus out-of-the-zone states. Furthermore, it is the fronto-parietal control network (FPCN) that drives this difference. Activity in the dorsal attention network (DAN) and DMN were desynchronized across both zone states. During in-the-zone periods, FPCN synchronized with DAN, while during out-of-the-zone periods, FPCN synchronized with DMN. In contrast, the ventral attention network synchronized more closely with DMN during in-the-zone periods compared to out-of-the-zone periods. These findings demonstrate that time-varying functional connectivity across different brain networks varies with fluctuations in sustained attention.

## Introduction

To successfully achieve one’s goals and perform optimally in many situations, we must control and sustain our attention. Tasks requiring such attentional control can vary from the mundane, like listening to a podcast, to highly engaging ones like playing basketball. Some think of attention as a light– it’s either on or off, but a better metaphor is a flickering candle–even when lit, the flame varies. These fluctuations of attention can occur from moment to moment and across more lengthy time periods (Mackworth, 1948; Dorrian et al., 2004; Esterman et al., 2013; Kucyi et al., 2017; Rosenberg et al., 2020). Sustaining attention is a complex cognitive process that requires both top-down (e.g., knowledge-driven processes to bias subject towards signal as opposed to noise) and bottom-up control (e.g., sensory inputs such as the characteristics of the target stimulus) (c.f., Sarter et al., 2001).

### Brain Networks of Sustained Attention

Scientists have worked to understand the neural mechanisms related to sustained attention (e.g., Mesulam, 1990; Dockree et al., 2004; Clayton et al., 2015; Rosenberg et al., 2016). From this work, one thing is clear, attention is not sustained through the activation of isolated brain regions, rather it is mediated by coordinated activity across multiple brain regions (c.f., Bressler & Menon, 2010). There are four major brain networks typically implicated in fluctuations in attention control (Yeo et al., 2011). They are the default mode (DMN), dorsal attention (DAN), ventral attention (VAN), and fronto-parietal control (FPCN) networks (Fortenbaugh et al., 2017; Esterman & Rothlein, 2019; Zuberer et al., 2021) (Figure 1).

**Figure 1:**
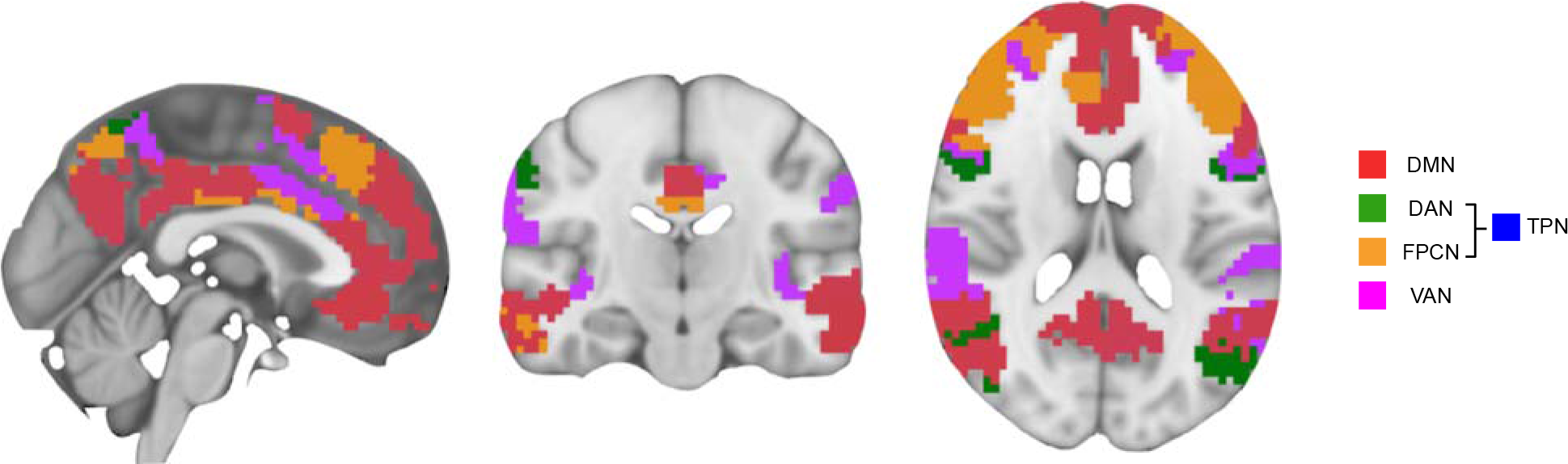
The Four Major Networks of the Brain Involved in Attention. Default mode network (DMN), dorsal attention network (DAN) and fronto-parietal control network (FPCN) combined is the task-positive network (TPN)(Petersen & Posner, 2012), and ventral attention network (VAN) (Schaefer et al., 2018).

The DAN is hypothesized to mediate the top-down task-oriented attention (Fox et al., 2005). The core brain regions are the frontal eye fields (FEF), superior parietal lobule (SPL), intraparietal sulcus (IPS), and precentral ventral frontal cortex (PrCv). The FPCN is believed to mediate executive control (Corbetta & Shulman, 2002; Vincent et al., 2008) and consists of the posterior dorsolateral prefrontal cortex (pDLPFC), the rostrolateral prefrontal cortex (RLPFC), anterior inferior parietal lobule (aIPL), posterior dorsomedial prefrontal cortex (pDMPFC), and middle temporal gyrus (MTG) (Yeo et al., 2011). Combined, these two networks are sometimes called the task-positive network (TPN) (Petersen & Posner, 2012).

The VAN is said to play a role in monitoring for salient inputs. It is sometimes referred to as the salience or ventral salience network (Menon & Uddin, 2010). The main cortical brain regions in the VAN are the anterior insula (AI) and dorsal anterior cingulate cortex (dACC). The model proposed by Bressler and Menon (2010) suggests that the antagonism between the DMN and DAN is mediated by the VAN. Further research has shown that the VAN works with FPCN and can couple with either DMN or DAN depending on whether one is attending to internal or externally directed goals, respectively (Vossel et al., 2014; Beaty et al., 2015).

Another association brain network implicated in task performance is the DMN. The core DMN brain regions include the medial prefrontal cortex (mPFC), the posterior cingulate cortex (PCC), precuneus, as well as the right frontal and left occipital regions. This network is most active during activities related to internal processes such as introspection, emotion perception, beliefs and intention, theory of mind, and mentalizing (Gusnard et al., 2001; Iacoboni et al., 2004; D’Argembeau et al., 2005; Spreng et al., 2009; Spreng & Grady, 2010). Furthermore, the DMN has frequently been found to deactivate during task performance and works in an antagonistic way with the TPN (Fox et al., 2005).

However, there is some disagreement in the literature on exactly how DMN activity supports behavior. Some studies suggest that the DMN is detrimental to performance (Weissman et al., 2006; Boly et al., 2007; Eichele et al., 2008; Christoff et al., 2009), while others claim that intermediate levels of DMN activity can improve outcomes (Gilbert et al., 2006; Hahn et al., 2007; Mason et al., 2007; Sadaghiani et al., 2009).

### Reaction Time Variability as a Behavioral Correlate of Sustained Attention

There may be many reasons for this ambiguity in the literature. A difficulty in studying sustained attention is finding a behavioral correlate to fluctuations of attention at shorter timescales. Classical vigilance tasks involve recognizing that a stimulus changes intermittently. This ability may fluctuate over minutes to hours (Mackworth, 1948; Dorrian et al., 2004), but given the low rate of required overt responses, some tasks cannot differentiate fluctuations that take place on faster timescales. More continuous tasks have been successfully used to measure attention at these timescales. For example, Conner’s CPT-II (Conners, 1994), gradCPT, a gradual change detection task (Rosenberg et al., 2013; Esterman et al., 2013), sustained attention to response task (SART) (Robertson et al.,1997), paced finger tapping task (Seli et al., 2013), self-paced finger tapping task (Kucyi et al., 2017), and breath counting task (Levinson et al., 2014) all have a high temporal sensitivity.

Additionally, while behavioral measures like error rates have been proposed to be good objective correlates to attention (Manly et al., 2000), Esterman and colleagues (2013) suggest that reaction time (RT) variability is a better trial-to-trial measure for studying fluctuating attentional states within an individual. Unusually slow RTs may indicate a lack of readiness or reduced attention to a task (Cheyne et al., 2009). Whereas abnormally fast RTs may indicate premature or routinized responding and have been associated with failures of attentional control and response inhibition (Weissman et al., 2006). Other studies (Bastian & Sackur, 2013; Seli et al., 2013) have also shown that deviations in performance variability correlate with mind wandering. In a recent experiment triangulating subjective experience with objective measures, Godwin and colleagues (2023) found that the highest average variance RT was reported when subjects subjectively judged themselves “off-task” and the lowest variance RT was reported when they thought they were “on-task.” Furthermore, intra-individual variability in RT has been linked to impairments of attention and executive function seen in attention-deficit hyperactivity disorder (Tamm et al., 2012), which supports the idea that erratic responding is related to greater deficits in attention. These results support the claim that RT variability can be used as an indicator of the level of sustained attention to a task.

Esterman and colleagues (2013) proposed categorizing behavior into two zone states based on RT variability. The first state, postulated to capture focused attention across time, is the in-the-zone state. It reflects optimal engagement with a task and is marked by stable responding, skillfulness, or mastery, that culminates in the perception of being in control. Esterman and colleagues speculate that in-the-zone captures the phenomenon *flow* (i.e., the engaged activity of capably performing a difficult task; Csikszentmihalyi, 1990). The second state, on the other hand, captures when attention wanes and we often feel out-of-the-zone. Being out-of-the-zone or colloquially, “zoned out”, is marked by an unstable performance that can lead to more errors. Suboptimal experiences can be on either extreme from under-engagement, capturing phenomena like boredom and mind wandering, to over-engagement, such as hyper-attentiveness due to overthinking (Esterman et al., 2014). Other reasons for feeling out-of-the-zone can be attributed to the lack of arousal or drowsiness which has a downstream effect on behavior (Godwin et al., 2023).

Esterman and colleagues (2013) found that sustained in-the-zone periods were associated with moderate DMN activity. Moreover, while in-the-zone, higher activity of DMN precedes and persists after an incorrect response, indicating that as automatic responding sets in, there may be a tendency to mind wander and in turn cause a lapse in attention. Conversely, when participants were out-of-the-zone there was less activity in DMN and higher activity in DAN. They posit that optimal performance may rely not just on activity in one network, but it might involve balancing activity between DMN and DAN. In the current study, we adopt the categorization of zone states to demarcate sustained attention performance as defined by Esterman and colleagues by RT variability.

### Time-varying Functional Connectivity to Capture Attention Fluctuations

Early studies of sustained attention using fMRI measured brain connectivity by computing regional correlations across the duration of the scan, which can last from seconds up to minutes (Adler et al., 2001; Lawrence et al., 2003; Strakowski et al., 2004). This assumption of stationarity is problematic given the fluctuating nature of attention. Additionally, researchers have found spatiotemporal activity and connectivity changes across seconds within a scan (Chang & Glover, 2010; Majeed et al., 2011; Liu & Dyun, 2013). Studies suggest that the time-varying properties of functional connectivity between regions can possibly produce different results depending on the timescale used to investigate the activity of the regions (Allen et al., 2012; Handwerker et al., 2012; Hutchison et al., 2012). These differences may explain the contradictory claims that DMN and DAN can both support and be detrimental to performance. Esterman and colleagues (2013) posit that a limitation may arise from looking at brain activation in isolation which does not provide a full neural mechanistic explanation for fluctuations of attention. Consequently, Kucyi and colleagues (2017) employed a time-varying measure of functional connectivity that tracks the variance of RT and the associated brain regions. They found that increased moment-to-moment RT variance correlates with increased functional connectivity between the DMN and VAN (also known as the salience network, SN, in their study).

Our study utilizes a different time-varying functional connectivity approach to bring clarity to the relationship between the association networks and how they relate to sustained attention.

### Framework of Sustained Attention

Kucyi and colleagues (2017) proposed a framework for sustained attention (shown in Figure 6a). During in-the-zone periods of sustained attention, they reported regional activation of the DMN and lower activity in the DAN and the VAN/SN. While out-of-the-zone periods were associated with lower activity of DMN and higher activity in the DAN and VAN. These results concur with the findings by Esterman and colleagues (2013). Furthermore, in-the-zone periods were marked by lower inter-regional functional connectivity within the regions of the DMN, and lower connectivity between DMN and salience network. While on the inverse, out-of-the-zone periods correlated with higher connectivity within DMN, and higher connectivity between DMN and salience.

Unfortunately, their framework focused on only the relationship between DMN and VAN. To attain a more complete understanding, our study investigates the inter-regional relationship between the attention networks (DAN, FPCN and VAN) and how they relate with the DMN. To do so, we employ a novel method that captures the repeating low-frequency fluctuations between the brain networks called the quasi-periodic pattern (QPP) analysis.

### Repeating Low-frequency Fluctuations in the Brain: The Quasi-periodic Pattern

In 1995, Biswal and colleagues noticed that there are neuronal fluctuations of <.01Hz with temporal coherence across the hemispheres of the brain and are larger in magnitude in the grey matter than the white matter. Consequently, this led many groups to interrogate these low-frequency fluctuations with varying methods on a variety of modalities. For example, there have been studies using electroencephalography (EEG) (Grooms et al., 2017; Leopold et al., 2003; Helfrich et al., 2018), local field potentials (LFPs) (Pan et al., 2013), and fMRI (Majeed et al., 2011; Thompson et al., 2014). Using EEG, researchers found that periodicity exists in the low frequency fluctuations (Valera et al., 1981; VanRullen & Koch, 2003) that may be linked to perception mainly through feedforward connections from sensory to association areas (Bastos et al., 2015, Spyropoulos et al., 2018). In 2018, Helfrich and colleagues supported this idea that the neural mechanism of sustained attention is rhythmic by showing attention-related theta-band (∼4Hz) oscillations of frontal and parietal cortical areas (regions of the FPCN) using intracranial EEG. Using fMRI, Majeed and colleagues (2011) reported spontaneous periodic repeating low frequency fluctuations that lasts approximately 20 s in humans. They called this the quasi-periodic pattern (QPP).

The QPP signal is marked by a reliably observed pattern of anti-correlation between the BOLD signal of the DMN and the TPN. Since then, the QPP has been found in humans during resting-state and task scans (Majeed et al., 2011; Thompson et al., 2013; Abbas et al., 2019a), in rats (Majeed et al., 2009; Majeed et al., 2011; Thompson et al., 2014), mice (Belloy et al., 2018), and macaques (Abbas et al., 2016) while awake or anesthetized. Of particular relevance to our study, in 2013, Thompson and colleagues discovered that faster RTs on a psychomotor vigilance task were significantly associated with higher anticorrelation of the DMN and the TPN. In a following study, they proposed that this anti-correlation pattern that they found is related to the QPP (Thompson et al., 2014).

Neural synchrony in the infraslow timescale may facilitate the coordination and organization of information processing in the brain (Buzsaki & Draguhn, 2004; Fox et al., 2005). This leads us to hypothesize that neural oscillations of in brain association networks, measured with methods like ones identifying the QPP, may capture fluctuations of attentional focus that occur on the order of seconds rather than minutes or longer (c.f., Raut et al., 2021). In the current study, we test this by investigating the relationship between the QPP and RT variability related to different zone states during the performance of a continuous finger tapping task. We hypothesize that the QPP will show greater segregation between DMN and TPN during low RT variability (viz., in-the-zone) when attention is sustained versus times of high RT variability (viz., out-of-the-zone) when attention lapses. Furthermore, with the QPP we can investigate how the subnetworks of the TPN (FPCN and DAN) and VAN relate to the DMN.

## Method

### Participants

Using the dataset from Godwin and colleagues (2023), there were 31 participants with fMRI scans and behavioral data. Their age ranged from 18 to 23 (M = 20, SD = 1.6). They were right-handed, had normal or corrected-to-normal vision, and did not report prior neurological or psychiatric conditions. To avoid including participants who may have fallen asleep, participants that tapped on less than 90% of the trials were excluded. This excluded two participants, leaving 29 participants (15 male, 13 female, and 1 gender-unidentified; average age: 19.6 ± 1.6 years) in this analysis.

### Task and Procedure

Participants performed a metronome response task (MRT) (c.f., Seli et al. 2013). They were instructed to tap along to a metronome tone as synchronously as possible. The task was organized into a series of blocks of tapping. These tapping blocks consisted of a 450-Hz tone presented for 75 ms. A 1300 ms of silence preceded each tone. In total, the metronome sounded at a rate of approximately .77 Hz (one tone per 1300 ms) (Figure 2). A baseline fixation cross of 2-4s preceded each tap period which remained on screen during the duration of the taps. There were five runs and 15 tapping blocks in each run which were made up of six tapping blocks of 16s, three blocks of 20s, two blocks of 24s, two blocks of 28s, one block of 32s, and one block of 36 s (run time = 10 mins and 33 seconds). The order of the blocks was randomized across runs.

**Figure 2:**
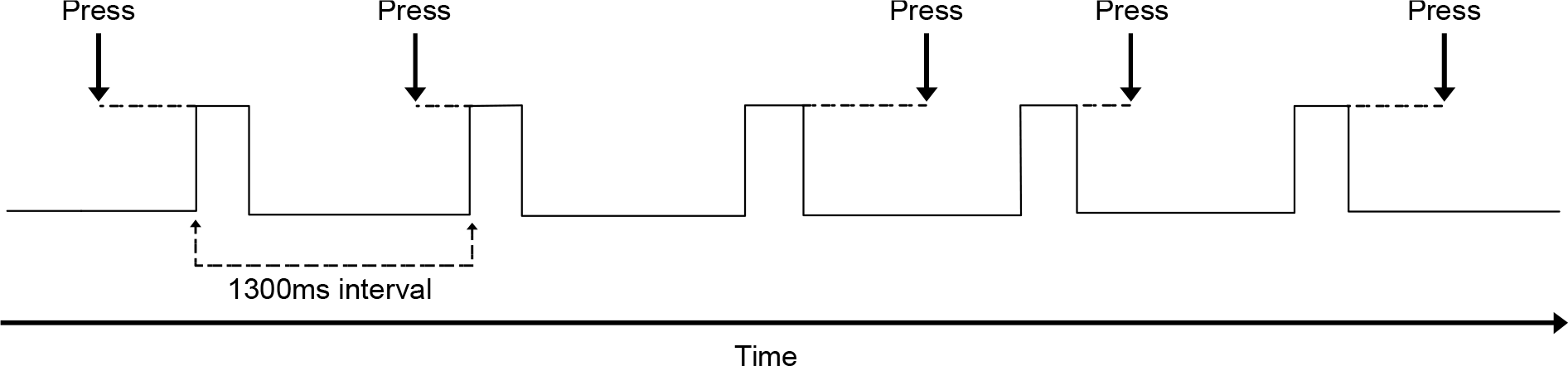
The Serial Tapping Task. Participants were instructed to tap along to a metronome tone as synchronously as possible. Each tone was separated by 1300 ms.

Participants tapped by pressing a button with their right index finger. After each tap period ended, thought probes were shown on the screen to measure their subjective rating of on-task and off-task. If participants selected the off-task option, they were then presented with two additional prompts to further address the nature of off-task thoughts. See Godwin and colleagues (2023) for the results of the subjective analysis.

### fMRI Design

Imaging was conducted on a Siemens 3T Trio MRI scanner at the GSU/GT Center for Advanced Brain Imaging. At the start, a T1-weighted MPRAGE anatomical scan was collected with the following acquisition parameters: FoV = 256 mm; 176 slices; 1.0 x 1.0 x 1.0 mm^3^ *voxels; flip angle = 9°*, TE = 3.98 ms; TR = 2250 ms; and TI = 850 ms. Next, participants completed the experiment consisted of five functional runs, each for a total duration of approximately 10 min and 33s. In particular, functional T2*-weighted echo-planar scans were collected during the runs with the following acquisition parameters: FoV = 204 mm; slices = 37; 3.0 x 3.0 x 3.0 mm^3^ voxels; interleaved slice acquisiton; gap = 0.5 mm; flip *angle = 90°*; TE = 30 ms; and TR = 2000 ms.

### Behavioral Data Analysis

Tapping RTs were analyzed using Seli and colleague’s method (2013) of calculating rhythmic response times (RRT) which is the difference in the time of a participant’s pressing of the button response box time-locked to the metronome’s tone onset time. RT variability for each run is calculated by taking the variance of each RRT within each run of the individual. Then a natural logarithm transformation was applied to adjust for the right-skewed distribution of RRT. Within each subject, the runs were rank ordered from highest to lowest RT variability. The run with the highest variability was labeled *out-of-the-zone* and the run with the lowest variability was labeled *in-the-zone* based on the categorization of Esterman and colleagues (2013).

### fMRI Data Preprocessing

Data preprocessing was performed using The Configurable Pipeline for the Analysis of Connectomes (C-PAC) (Craddock et al., 2013). This pipeline utilizes FMRIB Software Library (FSL) version 5.0 (Smith et al., 2004; Woolrich et al., 2009; Graham et al., 2016) and the Analysis of Functional NeuroImages (AFNI) software (Cox, 1996).

Anatomical scans (T1 images) were bias field corrected, skull stripped, and registered to the 2 mm Montreal Neurological Institute (MNI) atlas (Jenkinson & Smith, 2001; Jenkinson et al., 2002). Functional scans (EPI sequences) were slice-time and distortion corrected, masked, and motion corrected. Nuisance signal regression was done using the default settings of the C-PAC pipeline. Spatial smoothing was done using a Gaussian kernel with a full width at half maximum of 4 mm. Temporal filtering was set at a bandpass between 0.01 Hz and 0.1Hz. The top 5 principal components were calculated for white matter and cerebral spinal fluid using aCompCor and was extracted as nuisance regression. Global signal and cerebral spinal fluid mean signals were regressed after. Next, quadratic detrending was applied. All voxel time courses were z-scored to standardize the data for group-level analysis. The preprocessed images were then divided into the 7-network parcellation by Schaefer and colleagues (2018), which includes 400 defined regions of interest (ROI). The C-PAC pipeline is openly available at www.nitrc.org.

### Quasi-Periodic Pattern Template Acquisition and Analysis

A pattern-finding algorithm originally described by Majeed and colleagues (2011) and further refined (Yousefi & Keilholz, 2021; Xu et al., 2023) was applied separately to the concatenated brain sequences of all participant’s in-the-zone and out-of-the-zone runs. The algorithm starts with selecting a time window of 20s starting segment. Then the algorithm uses a sliding window correlation method to search for other instances where the BOLD signal in the brain sequence correlate with the starting segments’ peaks. As the search continues, additional segments are extracted and averaged, updating the segment. Notably, the search omits the final 20s of each concatenated run to avoid time boundary effects between different runs. This process repeats itself until the search can no longer find variations to the latest updated segment. By averaging similar segments, the output is a template of a reliable repeating pattern of activity within the functional scan also known as the QPP template. The QPP template is dominantly characterized by an anti-correlation between the DMN and TPN. The code for QPP analysis is openly available at https://github.com/GT-EmoryMINDlab/QPPLab.

The QPP template is represented as a normalized time-course of activity in each ROI of the parcellated brain regions. By taking the average of the sum of the ROIs in each network, we calculated a mean BOLD signal during each timepoint (TR) for the DMN and TPN within each zone state. We also looked at the subnetworks of TPN (DAN and FPCN) and VAN’s mean signal versus DMN within the QPP template. A positive correlation indicates that the networks behaved similarly across time, while a negative correlation signifies a decoupling of the networks.

## Results

### Statistical Analysis

To compare the correlation of the QPP timeseries between in-the-zone and out-of-the-zone, we compared the correlation coefficients of dependent groups with non-overlapping variables using the cocor package in R (Diedenhofen & Musch, 2015). It transforms the QPP correlation coefficients into Fisher’s *z*_r_ to make comparisons using the Pearson and Filon formula (1898, p. 262).

### DMN and TPN Correlation Comparison Within Zone States

The resulting QPP template in the DMN and TPN in each zone state is shown in Figure 3. Pairwise correlations of the QPP template for DMN and TPN of in-the-zone (*r* = −0.94) and out-of-the zone (*r* = +0.67) were shown to be significantly different across the timecourse, *z* = −11.83, *p* < .001.

**Figure 3:**
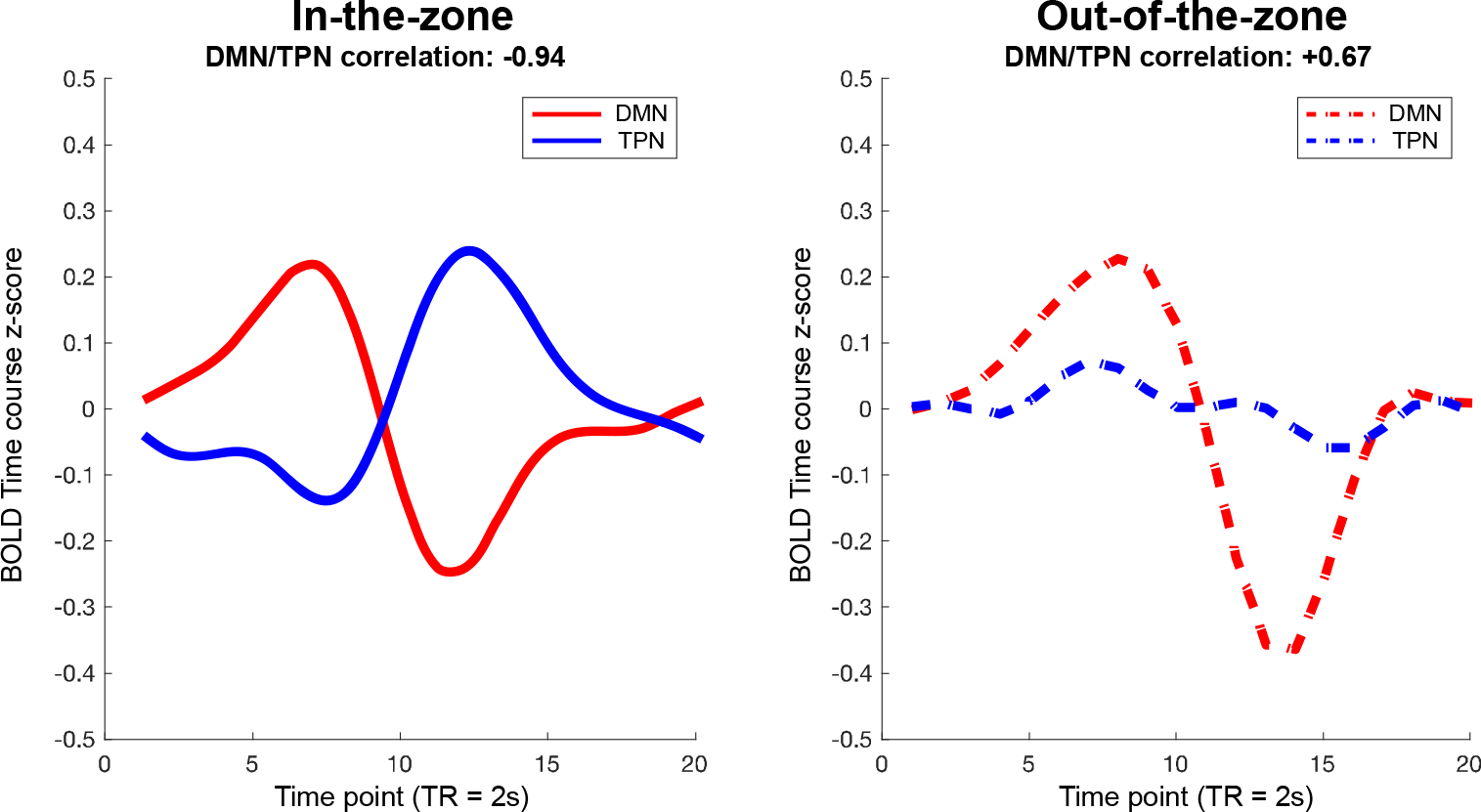
The QPP template of DMN and TPN. In-the-zone has a negative correlation of −0.94 while out-of-the-zone has a positive correlation of +0.67 and they are statistically different (p<.05)

### DMN’s Relationship With DAN, FPCN, and VAN Within Zone States

To further interrogate the relationship of other networks hypothesized to play a role in sustained attention, we plotted all the subnetworks of TPN, and the VAN. As shown in Figure 4, DAN activity was unaffected by zone state, *z* = −1.95, *p* > .05, negatively correlating with DMN in both in-the-zone (*r* = - 0.99) and out-of-the-zone (*r* = −0.95). On the other hand, the FPCN decoupled from DMN while in-the-zone (*r* = −0.35) and correlated with DMN while subjects were out-of-the-zone (*r* = +0.98), *z* = −6.70, *p*< .001. VAN showed the opposite pattern to FPCN. It was positively correlated with DMN during in-the-zone (*r* = +0.33) and negatively correlated with DMN during out-of-the-zone (*r* = −0.97), *z* = 6.36, *p* < .001.

**Figure 4:**
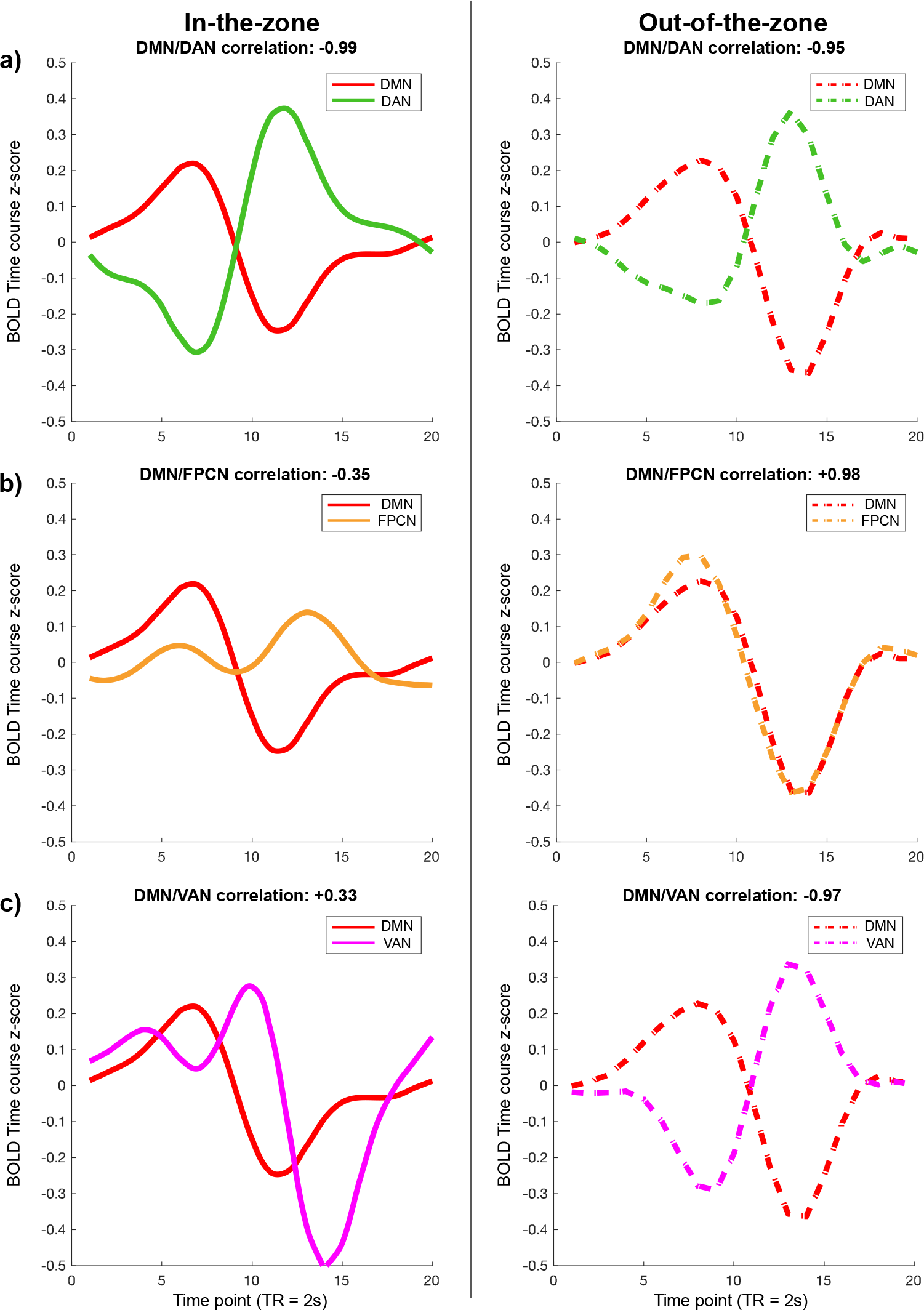
Plots of Each of the Attention Networks in Relation to DMN (a) DAN is not differentiated between the networks but is largely responsible for driving the anti-correlation of the QPP template (b) FPCN is less segregated from DMN in out-of-the-zone compared to in-the-zone (c) VAN is less segregated from DMN in in-the-zone compared to out-of-the-zone.

### Comparison of Networks Across Zone States

To further investigate how fluctuations in attention affected the QPP within networks, we compared each network across zone states (Figure 5). The DMN was significantly positively correlated between zones, *r* = +0.66, *t*(18) = 3.75, *p* < .01. Similarly, DAN was also significantly positively correlated at, *r* =+0.80, *t*(18) = 5.67, *p* < .001. Whereas activity in FPCN during each zone state was significantly negatively correlated, *r* = −0.59, *t*(18) = −3.09, *p* < .01. Lastly, the VAN was significantly different across conditions, *r* = −0.83, *t*(18) = −6.33, *p* < .001.

**Figure 5:**
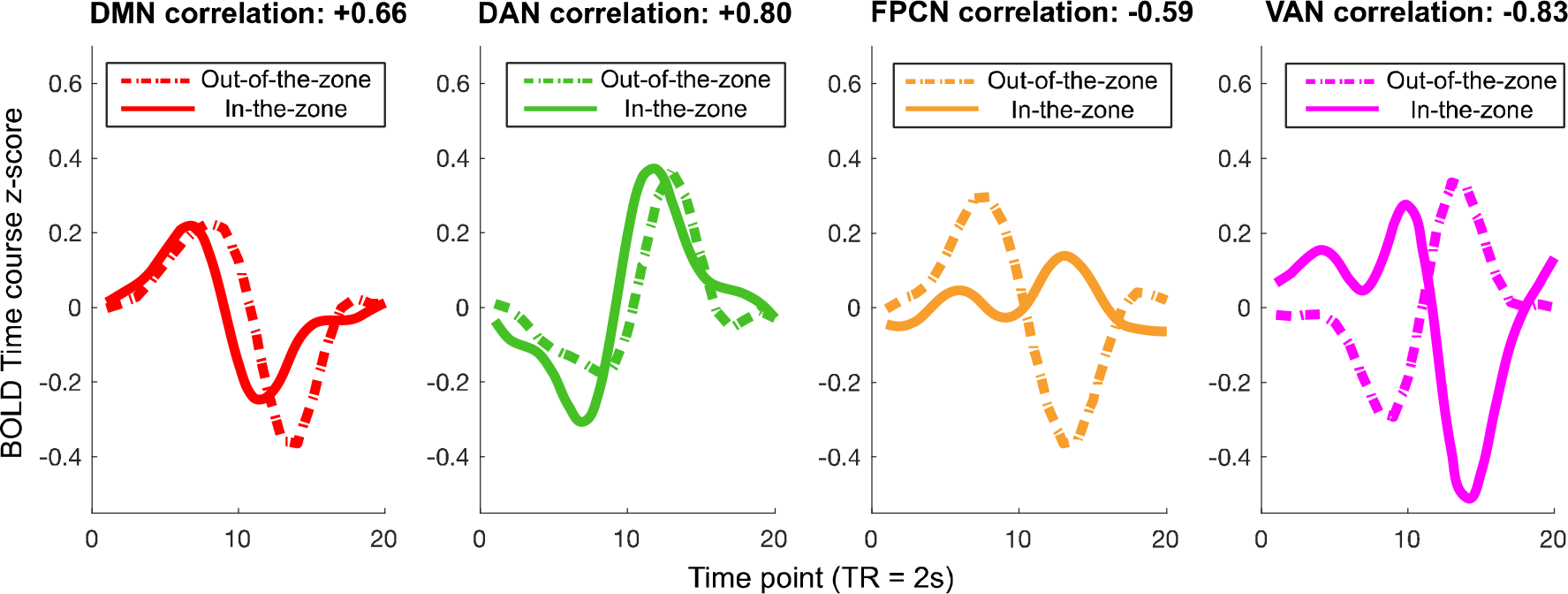
The Associations Networks Compared Between Conditions. Both DMN and DAN reported a positive correlation denoting similar phase and magnitude. The FPCN in both zone states were negatively correlated which drove the difference between the DMN/TPN relationship. The VAN reported the most difference between the conditions due to a flip in negative and positive phase.

## Discussion

In this study, we investigated the relationship between brain networks during moments of more or less engaged (i.e., sustained) attention. Previous research has shown changes in network relationships relating to sustained attention (Esterman et al., 2013; Kucyi et al., 2017). The current study investigated dynamic changes in network connectivity as attention fluctuates. Specifically, we interrogated low frequency fluctuations, commonly called QPPs, and hypothesized that QPP relationship between DMN and TPN would differ in in-the-zone and out-of-the-zone performance. This hypothesis was supported (Figure 3). Additionally, we discovered that this change was primarily driven by attention-related changes in the FPCN. We also found that VAN activity differentiated attentional zone states (Figure 4 & 5).

### Sustained Attention in DMN, TPN, and TPN Subnetworks (FPCN & DAN)

As predicted, the QPP relationship between DMN and TPN during the in-the-zone condition was significantly more anti-correlated than out-of-the-zone’s (Figure 5). That is, more successful sustained attention was associated with more segregation between these networks. This result complements prior research showing that faster response is associated with higher anti-correlation between DMN and TPN (Thompson et al., 2013). Moreover, Abbas and colleagues (2019b) found a similar relationship in ADHD patients. Specifically, they found that the QPP was more segregated in healthy controls than in patients with a chronic impairment of sustained attention. Taken together, the results suggest that the decoupling of the two networks in the low frequency brain activity is correlated with better sustained attention. This antagonistic relationship between the networks has also been reported in many studies using static connectivity (Kelley et al., 2008; Hoekzema et al., 2013; Magnuson et al., 2015).

We found that the subnetworks within TPN were differentially related to zone states. Changes in FPCN connectivity across conditions drove the DMN-TPN differences between in-the-zone and out-of-the-zone states, whereas DAN and DMN’s relationship remained similar across the states (Figure 4a & 4b). Further comparison of the FPCN from each zone state showed that they were significantly negatively correlated with each other (Figure 5). This reaffirms that what drives the significant difference between DMN and TPN correlation of the zone states stems from the FPCN. Previous research has shown that DAN and FPCN have different relationships with DMN (Yousefi & Keilholz, 2021), but this is the first demonstration that these relationships are related to sustained attention and suggests that the FPCN plays an import role in engaged attention– at least in this task.

Although Petersen and Posner (2012) identified the regions of the FPCN as part of the TPN, our results suggest that the role this network plays in task performance may depend on an individuals current attentional state. During engaged attentional states like in-the-zone, FPCN does indeed synchronize with DAN, resulting in the canonical TPN identified by Petersen and Posner (2012). However, during out-of-the-zone periods, FPCN synchronizes instead with DMN and may be less appropriately characterized as “task positive.”

Research suggests a possible mechanistic explanation for FPCN that may clarify this change in activation pattern during different zone states. Unsworth and Robison (2017) propose that FPCN activity suppresses DMN during externally demanding attention tasks. Further support for this idea comes from Spreng and colleagues (2013) who found evidence of FPCN mediating internal and external goals by flexibly coupling with DMN or DAN. These ideas are consistent with our data, FPCN may support sustained attention during in-the-zone state by synchronizing with an externally oriented network (DAN) or it may hamper sustained attention during out-of-the-zone states by synchronizing with an internally oriented network (DMN).

### Sustained Attention and Brain Network Connectivity

Interestingly, as opposed to the FPCN, the VAN was more anti-correlated with DMN in out-of-the-zone states rather than in-the-zone states (Figure 4c). Recent interrogation of the QPP in resting state finds that DMN tends to anti-correlate with both DAN and VAN (Yousefi & Keilholz, 2021). Hence, it is not surprising that VAN and DMN pattern in out-of-the-zone state matches the QPP found in resting state scans as the out-of-the-zone state seems more closely related to resting state brain processing than in-the-zone. What is peculiar is that VAN synchronizes more closely with DMN in in-the-zone than out-of-the-zone states. More task based QPP research is needed to understand why that might be the case.

Our study adds to the framework of sustained-attention proposed by Kucyi and colleagues (2017) (Figure 6b) by using a novel time-varying functional connectivity method of the QPP analysis to understand the low frequency fluctuation fluctuations correlated with sustained attention.

**Figure 6:**
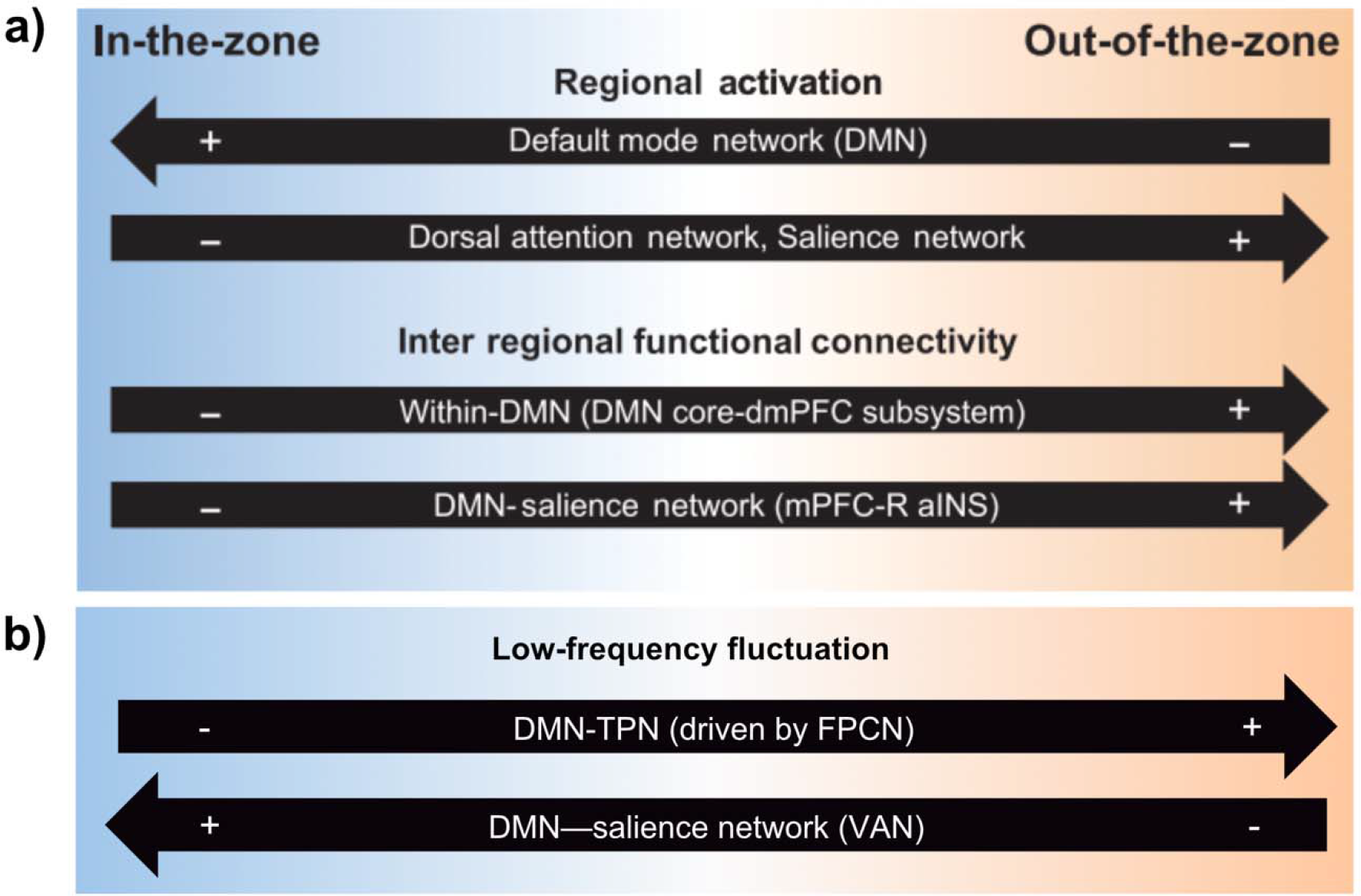
Framework of Sustained Attention During Zone State Performance (a) for regional activation and inter regional functional connectivity framework (Kucyi et al., 2017). (b) low frequency fluctuation framework from our results.

### Discrepancy and Limitations

In our study, we found that VAN (salience network) was positively correlated with DMN when participants were in-the-zone, and negatively correlated with DMN when they were out-of-the-zone. This contradicts the results from Kucyi and colleagues (2017) that reported the opposite relationship using static (cross-run) measures of connectivity. Research shows that QPP contributes significantly to the BOLD signal across runs (Abbas et al., 2019a; Abbas et al., 2019b), so it is surprising that the current results do not align with Kucyi and colleages. There may be multiple reasons for this discrepancy. First, there are many differences in the analyses. We removed global signal while Kucyi and colleagues did not report conducting global signal regression. Presence or absence of global signal is known to substantially alter functional connectivity (Murphy et al., 2009; Murphy & Fox, 2017; Yousefi et al., 2018).

Furthermore, our signal was band-pass filtered between 0.01 Hz and 0.1Hz as opposed the 0.01 Hz high pass that they applied to their data. Low-frequency fluctuations have been suggested to modulate long-distance neuronal synchronization while high-frequency fluctuations are thought as localized regional network activity, hence the lack of representation of higher frequencies in our data could present divergent results (Von Stein & Sarnthein, 2000; Müller et al., 2011; Siegel et al., 2012). The different temporal frequencies may reflect different signaling along the same anatomical pathways (Helfrich & Knight, 2016). Finally, we used a parcellation based method to identify mean signal across all ROIs in each network. In contrast, Kucyi and colleagues used a seed-based approach. We attempted to rule out this last difference by taking the defined ROIs from their study of right mPFC (xyz = 6, 66, 4) and right anterior insula (xyz = 40, 8, 0) and matched them with the closest parcel in Schaefer’s 400roi parcellation. We then computed group functional connectivity in just these parcels. The results were more consistent with our QPP results (and inconsistent with Kucyi et al., 2017) where functional connectivity between the two regions was more negative in out-of-the-zone than in-the-zone (X and Y, respectively), but the minor difference was not significant and fails to resolve the discrepancy. Thus, it may be the specific patterns of DMN and VAN connectivity depend on other differences between the experiments, or the pattern is less reliable than previously proposed.

### Conclusion

This study is the first to employ low-frequency fluctuations with a network-based approach to understand the neural mechanisms of sustained-attention intra-individually. We found that the FPCN is important in integrating with DAN and disassociating with DMN for in-the-zone performance. Secondly, VAN works coherently with DMN during in-the-zone states in contrast with out-of-the-zone states. These results begin to identify the complex role these networks play in mediating attention across short time scales. More work is necessary to clarify the mechanisms for how these dynamic changes in activity and connectivity relate to the static changes previously described in the literature (e.g., investigating how preprocessing and processing strategies can affect the results) to mediate fluctuations in sustained attention.

### Open Practices Statement

Data or materials for the experiments are available upon request, while scripts related to the QPP algorithm are openly available at https://github.com/GT-EmoryMINDlab/QPPLab. Certain parts of the analyses were preregistered at https://osf.io/tcvgd. The rest of the preregistered analyses are beyond the scope of this paper and have yet to be completed.

## References

Abbas, A., Langley, J., Howell, L., & Keilholz, S. (2016). Quasiperiodic patterns vary in frequency between anesthetized and awake monkeys. In Resting state brain connectivity biennial conference (p. 141).

Abbas, A., Belloy, M., Kashyap, A., Billings, J., Nezafati, M., Schumacher, E. H., & Keilholz, S. (2019a). Quasi-periodic patterns contribute to functional connectivity in the brain. Neuroimage, 191, 193–204. https://doi.org/10.1016/j.neuroimage.2019.01.076

Abbas, A., Bassil, Y., & Keilholz, S. D. (2019). Quasi-periodic patterns of brain activity in individuals with attention-deficit/hyperactivity disorder. NeuroImage: Clinical, 21, 101653. https://doi.org/10.1016/j.nicl.2019.101653

Adler, C. M., Sax, K. W., Holland, S. K., Schmithorst, V., Rosenberg, L., & Strakowski, S. M. (2001). Changes in neuronal activation with increasing attention demand in healthy volunteers: An fMRI study. Synapse, 42(4), 266–272. https://doi.org/10.1002/syn.1112

Allen, E., Damaraju, E., Plis, S. M., Erhardt, E. B., Eichele, T., & Calhoun, V. D. (2014). Tracking Whole-Brain Connectivity Dynamics in the Resting State. Cerebral Cortex, 24(3), 663–676. https://doi.org/10.1093/cercor/bhs352

Bastian, M., & Sackur, J. (2013). Mind wandering at the fingertips: automatic parsing of subjective states based on response time variability. Frontiers in Psychology, 4. https://doi.org/10.3389/fpsyg.2013.00573

Bastos, A. M., Vezoli, J., Bosman, C. A., Schoffelen, J., Oostenveld, R., Dowdall, J. R., De Weerd, P., Kennedy, H., & Fries, P. (2015). Visual Areas Exert Feedforward and Feedback Influences through Distinct Frequency Channels. Neuron, 85(2), 390–401. https://doi.org/10.1016/j.neuron.2014.12.018

Beaty, R. E., Holling, H., Kaufman, S. B., & Silvia, P. J. (2015). Default and Executive Network Coupling Supports Creative Idea Production. Scientific Reports, 5(1). https://doi.org/10.1038/srep10964

Belloy, M. E., Naeyaert, M., Abbas, A., Shah, D., Vanreusel, V., Van Audekerke, J., Keilholz, S. D., Keliris, G. A., Van Der Linden, A., & Verhoye, M. (2018). Dynamic resting state fMRI analysis in mice reveals a set of Quasi-Periodic Patterns and illustrates their relationship with the global signal. NeuroImage, 180, 463–484. https://doi.org/10.1016/j.neuroimage.2018.01.075

Biswal, B. B., Yetkin, F. Z., Haughton, V. M., & Hyde, J. S. (1995). Functional connectivity in the motor cortex of resting human brain using echo-planar mri. Magnetic Resonance in Medicine, 34(4), 537–541. https://doi.org/10.1002/mrm.1910340409

Boly, M., Balteau, E., Schnakers, C., Degueldre, C., Moonen, G., Luxen, A., Phillips, C., Peigneux, P., Maquet, P., & Laureys, S. (2007). Baseline brain activity fluctuations predict somatosensory perception in humans. Proceedings of the National Academy of Sciences of the United States of America, 104(29), 12187–12192. https://doi.org/10.1073/pnas.0611404104

Bressler, S. L., & Menon, V. (2010). Large-scale brain networks in cognition: emerging methods and principles. Trends in Cognitive Sciences, 14(6), 277–290. https://doi.org/10.1016/j.tics.2010.04.004

Buzsáki, G., & Draguhn, A. (2004). Neuronal Oscillations in Cortical Networks. Science, 304(5679), 1926– 1929. https://doi.org/10.1126/science.1099745

Chang, C., & Glover, G. H. (2010). Time–frequency dynamics of resting-state brain connectivity measured with fMRI. NeuroImage, 50(1), 81–98. https://doi.org/10.1016/j.neuroimage.2009.12.011

Cheyne, J. A., Solman, G. J. F., Carriere, J. S. A., & Smilek, D. (2009). Anatomy of an error: A bidirectional state model of task engagement/disengagement and attention-related errors. Cognition, 111(1), 98–113. https://doi.org/10.1016/j.cognition.2008.12.009

Christoff, K., Irving, Z. C., Fox, K. C. R., Spreng, R. N., & Andrews-Hanna, J. R. (2016). Mind-wandering as spontaneous thought: a dynamic framework. Nature Reviews Neuroscience, 17(11), 718–731. https://doi.org/10.1038/nrn.2016.113

Clayton, M. C., Yeung, N., & Kadosh, R. C. (2015). The roles of cortical oscillations in sustained attention. Trends in Cognitive Sciences, 19(4), 188–195. https://doi.org/10.1016/j.tics.2015.02.004

Conners, C. K. (2000). Conners’ continuous performance test. Multi-Health Systems.

Corbetta, M., & Shulman, G. L. (2002). Control of goal-directed and stimulus-driven attention in the brain. Nature Reviews Neuroscience, 3(3), 201–215. https://doi.org/10.1038/nrn755

Cox, R. A. (1996). AFNI: Software for Analysis and Visualization of Functional Magnetic Resonance Neuroimages. Computers and Biomedical Research, 29(3), 162–173. https://doi.org/10.1006/cbmr.1996.0014

Craddock, R. C., James, G. A., Holtzheimer, P. E., Hu, X., & Mayberg, H. S. (2012). A whole brain fMRI atlas generated via spatially constrained spectral clustering. Human Brain Mapping, 33(8), 1914– 1928. https://doi.org/10.1002/hbm.21333

Csikszentmihalyi, M. (2009). Flow: The Psychology of Optimal Experience. Harper Collins.

D’Argembeau, A., Collette, F., Van Der Linden, M., Laureys, S., Del Fiore, G., Degueldre, C., Luxen, A., & Salmon, E. (2005). Self-referential reflective activity and its relationship with rest: a PET study. NeuroImage, 25(2), 616–624. https://doi.org/10.1016/j.neuroimage.2004.11.048

Diedenhofen, B. & Musch, J. (2015). cocor: A Comprehensive Solution for the Statistical Comparison of Correlations. PLoS ONE, 10(4): e0121945. https://doi:10.1371/journal.pone.0121945

Dockree, P. M., Kelly, S. P., Roche, R. a. P., Hogan, M., Reilly, R. B., & Robertson, I. H. (2004). Behavioural and physiological impairments of sustained attention after traumatic brain injury. Cognitive Brain Research, 20(3), 403–414. https://doi.org/10.1016/j.cogbrainres.2004.03.019

Dorrian, J., Rogers, N. L., & Dinges, D. F. (2004). Psychomotor Vigilance Performance: Neurocognitive Assay Sensitive to Sleep Loss. CRC Press EBooks, 39–70. https://doi.org/10.3109/9780203998007-4

Eichele, T., Debener, S., Calhoun, V. D., Specht, K., Engel, A., Hugdahl, K., Von Cramon, D. Y., & Ullsperger, M. (2008). Prediction of human errors by maladaptive changes in event-related brain networks. Proceedings of the National Academy of Sciences of the United States of America, 105(16), 6173–6178. https://doi.org/10.1073/pnas.0708965105

Esterman, M., & Rothlein, D. (2019). Models of sustained attention. Current Opinion in Psychology, 29, 174–180. https://doi.org/10.1016/j.copsyc.2019.03.005

Esterman, M., Noonan, S., Hedeker, D., & DeGutis, J. (2013). In the Zone or Zoning Out? Tracking Behavioral and Neural Fluctuations During Sustained Attention. Cerebral Cortex, 23(11), 2712– 2723. https://doi.org/10.1093/cercor/bhs261

Esterman, M., Rosenberg, M. D., & Noonan, S. K. (2014). Intrinsic fluctuations in sustained attention and distractor processing. Journal of Neuroscience, 34(5), 1724–1730. https://doi.org/10.1523/jneurosci.2658-13.2014

Fortenbaugh, F. C., DeGutis, J., & Esterman, M. (2017). Recent theoretical, neural, and clinical advances in sustained attention research. Annals of the New York Academy of Sciences, 1396(1), 70–91. https://doi.org/10.1111/nyas.13318

Fox, M. D., Snyder, A. Z., Vincent, J. L., Corbetta, M., Van Essen, D. C., & Raichle, M. E. (2005). The human brain is intrinsically organized into dynamic, anticorrelated functional networks. Proceedings of the National Academy of Sciences of the United States of America, 102(27), 9673–9678. https://doi.org/10.1073/pnas.0504136102

Gilbert, S. J., Simons, J. S., Frith, C. D., & Burgess, P. W. (2006). Performance-related activity in medial rostral prefrontal cortex (area 10) during low-demand tasks. Journal of Experimental Psychology: Human Perception and Performance, 32(1), 45–58. https://doi.org/10.1037/0096-1523.32.1.45

Godwin, C. A., Smith, D. M., & Schumacher, E. H. (2023). Beyond mind wandering: Performance variability and neural activity during off-task thought and other attention lapses. Consciousness and Cognition, 108, 103459. https://doi.org/10.1016/j.concog.2022.103459

Graham M, Drobnjak I, Zhang H, editors. Quantitative evaluation of eddy-current motion correction techniques for diffusion-weighted MRI. In: International Society for Magnetic Resonance in Medicine (ISMRM). Singapore (2016).

Grooms, J. K., Thompson, G. J., Pan, W., Billings, J., Schumacher, E. H., Epstein, C. J., & Keilholz, S. D. (2017). Infraslow Electroencephalographic and Dynamic Resting State Network Activity. Brain Connectivity, 7(5), 265–280. https://doi.org/10.1089/brain.2017.0492

Gusnard, D. A., Akbudak, E., Shulman, G. L., & Raichle, M. E. (2001). Medial prefrontal cortex and self-referential mental activity: Relation to a default mode of brain function. Proceedings of the National Academy of Sciences of the United States of America, 98(7), 4259–4264. https://doi.org/10.1073/pnas.071043098

Hahn, B., Ross, T. W., & Stein, E. A. (2007). Cingulate Activation Increases Dynamically with Response Speed under Stimulus Unpredictability. Cerebral Cortex, 17(7), 1664–1671. https://doi.org/10.1093/cercor/bhl075

Handwerker, D. A., Roopchansingh, V., Gonzalez-Castillo, J., & Bandettini, P. A. (2012). Periodic changes in fMRI connectivity. NeuroImage, 63(3), 1712–1719. https://doi.org/10.1016/j.neuroimage.2012.06.078

Helfrich, R. F., & Knight, R. T. (2016). Oscillatory Dynamics of Prefrontal Cognitive Control. Trends in Cognitive Sciences, 20(12), 916–930. https://doi.org/10.1016/j.tics.2016.09.007

Helfrich, R. F., Fiebelkorn, I. C., Szczepanski, S. M., Lin, J., Parvizi, J., Knight, R. T., & Kastner, S. (2018b). Neural Mechanisms of Sustained Attention Are Rhythmic. Neuron, 99(4), 854–865.e5. https://doi.org/10.1016/j.neuron.2018.07.032

Hoekzema, E., Carmona, S., Ramos-Quiroga, J. A., Fernández, V. R., Bosch, R., Soliva, J. C., Rovira, M., Bulbena, A., Tobeña, A., Casas, M., & Vilarroya, O. (2014). An independent components and functional connectivity analysis of resting state fMRI data points to neural network dysregulation in adult ADHD. Human Brain Mapping, 35(4), 1261–1272. https://doi.org/10.1002/hbm.22250

Hutchison, R. M., Womelsdorf, T., Gati, J. S., Everling, S., & Menon, R. S. (2013). Resting-state networks show dynamic functional connectivity in awake humans and anesthetized macaques. Human Brain Mapping, 34(9), 2154–2177. https://doi.org/10.1002/hbm.22058

Iacoboni, M., Lieberman, M. D., Knowlton, B. J., Molnar-Szakacs, I., Moritz, M., Throop, C. J., & Fiske, A. P. (2004). Watching social interactions produces dorsomedial prefrontal and medial parietal BOLD fMRI signal increases compared to a resting baseline. NeuroImage, 21(3), 1167– 1173. https://doi.org/10.1016/j.neuroimage.2003.11.013

Jenkinson, M., & Smith, S. M. (2001). A global optimisation method for robust affine registration of brain images. Medical Image Analysis, 5(2), 143–156. https://doi.org/10.1016/s1361-8415(01)00036-6

Jenkinson, M., Bannister, P., Brady, M. E., & Smith, S. M. (2002). Improved Optimization for the Robust and Accurate Linear Registration and Motion Correction of Brain Images. NeuroImage, 17(2), 825–841. https://doi.org/10.1006/nimg.2002.1132

Kelly, A. M. C., Uddin, L. Q., Biswal, B. B., Castellanos, F. X., & Milham, M. P. (2008). Competition between functional brain networks mediates behavioral variability. NeuroImage, 39(1), 527–537. https://doi.org/10.1016/j.neuroimage.2007.08.008

Kucyi, A., Hove, M. J., Esterman, M., Hutchison, R. M., & Valera, E. M. (2017). Dynamic brain network correlates of spontaneous fluctuations in attention. Cerebral cortex, 27(3), 1831–1840.

Lawrence, N., Ross, T. W., Hoffmann, R. G., Garavan, H., & Stein, E. A. (2003). Multiple Neuronal Networks Mediate Sustained Attention. Journal of Cognitive Neuroscience, 15(7), 1028–1038. https://doi.org/10.1162/089892903770007416

Leopold, D. A., Murayama, Y., & Logothetis, N. K. (2003). Very Slow Activity Fluctuations in Monkey Visual Cortex: Implications for Functional Brain Imaging. Cerebral Cortex, 13(4), 422–433. https://doi.org/10.1093/cercor/13.4.422

Levinson, D. M., Stoll, E. L., Kindy, S., Merry, H. L., & Davidson, R. J. (2014). A mind you can count on: validating breath counting as a behavioral measure of mindfulness. Frontiers in Psychology, 5. https://doi.org/10.3389/fpsyg.2014.01202

Liu, X., & Duyn, J. H. (2013). Time-varying functional network information extracted from brief instances of spontaneous brain activity. Proceedings of the National Academy of Sciences of the United States of America, 110(11), 4392–4397. https://doi.org/10.1073/pnas.1216856110

Mackworth, N. H. (1948). The Breakdown of Vigilance during Prolonged Visual Search. Quarterly Journal of Experimental Psychology, 1(1), 6–21. https://doi.org/10.1080/17470214808416738

Magnuson, M. L., Thompson, G. J., Schwarb, H., Pan, W., Mckinley, A., Schumacher, E. H., & Keilholz, S. D. (2015). Errors on interrupter tasks presented during spatial and verbal working memory performance are linearly linked to large-scale functional network connectivity in high temporal resolution resting state fMRI. Brain Imaging and Behavior. https://doi.org/10.1007/s11682-014-9347-3

Majeed, W., Magnuson, M. L., & Keilholz, S. D. (2009). Spatiotemporal dynamics of low frequency fluctuations in BOLD fMRI of the rat. Journal of Magnetic Resonance Imaging, 30(2), 384–393. https://doi.org/10.1002/jmri.21848

Majeed, W., Magnuson, M. L., Hasenkamp, W., Schwarb, H., Schumacher, E. H., Barsalou, L. W., & Keilholz, S. D. (2011). Spatiotemporal dynamics of low frequency BOLD fluctuations in rats and humans. NeuroImage, 54(2), 1140–1150. https://doi.org/10.1016/j.neuroimage.2010.08.030

Manly, T., Davison, B., Heutink, J., Galloway, M., & Robertson, I. H. (2000). Not enough time or not enough attention? Speed, error and self-maintained control in the Sustained Attention to Response Test (SART). Clinical Neuropsychological Assessment!Z: An International Journal for Research & Clinical Practice, 3, 167–177.

Mason, M. F., Norton, M. I., Van Horn, J. D., Wegner, D. M., Grafton, S. T., & Macrae, C. N. (2007). Wandering Minds: The Default Network and Stimulus-Independent Thought. Science, 315(5810), 393–395. https://doi.org/10.1126/science.1131295

Menon, V., & Uddin, L. Q. (2010). Saliency, switching, attention and control: a network model of insula function. Brain Structure & Function, 214(5–6), 655–667. https://doi.org/10.1007/s00429-010-0262-0

Mesulam, M. (1990). Large-scale neurocognitive networks and distributed processing for attention, language, and memory. Annals of Neurology, 28(5), 597–613. https://doi.org/10.1002/ana.410280502

Müller, R., Shih, P., Keehn, B., Deyoe, J. R., Leyden, K. M., & Shukla, D. K. (2011). Underconnected, but How? A Survey of Functional Connectivity MRI Studies in Autism Spectrum Disorders. Cerebral Cortex, 21(10), 2233–2243. https://doi.org/10.1093/cercor/bhq296

Murphy, K., & Fox, M. D. (2017). Towards a consensus regarding global signal regression for resting state functional connectivity MRI. NeuroImage, 154, 169–173. https://doi.org/10.1016/j.neuroimage.2016.11.052

Murphy, K., Birn, R. M., Handwerker, D. A., Jones, T. R., & Bandettini, P. A. (2009). The impact of global signal regression on resting state correlations: Are anti-correlated networks introduced? NeuroImage, 44(3), 893–905. https://doi.org/10.1016/j.neuroimage.2008.09.036

Padilla, M., Wood, R. D., Hale, L. P., & Knight, R. T. (2006). Lapses in a Prefrontal-Extrastriate Preparatory Attention Network Predict Mistakes. Journal of Cognitive Neuroscience, 18(9), 1477–1487. https://doi.org/10.1162/jocn.2006.18.9.1477

Pan, W., Thompson, G. J., Magnuson, M. L., Jaeger, D., & Keilholz, S. D. (2013b). Infraslow LFP correlates to resting-state fMRI BOLD signals. NeuroImage, 74, 288–297. https://doi.org/10.1016/j.neuroimage.2013.02.035

Pearson, K., & Filon, L. N. G. (1898). Mathematical contributions to theory of evolution: IV. On the probable errors of frequency constants and on the influence of random selection and correlation. Philosophical Transactions of the Royal Society of London, Series A, 191, 229–311. doi:10.1098/rsta.1898.0007

Petersen, S. E., & Posner, M. I. (2012). The Attention System of the Human Brain: 20 Years After. Annual Review of Neuroscience, 35(1), 73–89. https://doi.org/10.1146/annurev-neuro-062111-150525

Raut, R. V., Snyder, A. Z., Mitra, A., Yellin, D., Fujii, N., Malach, R., & Raichle, M. E. (2021). Global waves synchronize the brain’s functional systems with fluctuating arousal. Science Advances, 7(30). https://doi.org/10.1126/sciadv.abf2709

Robertson, I. H., Manly, T., Andrade, J., Baddeley, B., & Yiend, J. (1997). Performance correlates of everyday attentional failures in traumatic brain injured and normal subjects. Neuropsychologia, 35(6), 747–758. https://doi.org/10.1016/s0028-3932(97)00015-8

Rosenberg, M. D., Finn, E. S., Scheinost, D., Papademetris, X., Shen, X., Constable, R. T., & Chun, M. M. (2016). A neuromarker of sustained attention from whole-brain functional connectivity. Nature neuroscience, 19(1), 165–171. https://doi.org/10.1038/nn.4179

Rosenberg, M., Noonan, S., DeGutis, J., & Esterman, M. (2013). Sustaining visual attention in the face of distraction: a novel gradual-onset continuous performance task. Attention, Perception, & Psychophysics, 75(3), 426–439. https://doi.org/10.3758/s13414-012-0413-x

Rosenberg, M. D., Scheinost, D., Greene, A. S., Avery, E. W., Kwon, Y. H., Finn, E. S., Ramachandran, R., Qiu, M., Constable, R. T., & Chun, M. M. (2020). Functional connectivity predicts changes in attention observed across minutes, days, and months. Proceedings of the National Academy of Sciences, 117(7), 3797–3807. https://doi.org/10.1073/pnas.1912226117

Sadaghiani, S., Hesselmann, G., & Kleinschmidt, A. (2009). Distributed and Antagonistic Contributions of Ongoing Activity Fluctuations to Auditory Stimulus Detection. The Journal of Neuroscience, 29(42), 13410–13417. https://doi.org/10.1523/jneurosci.2592-09.2009

Sarter, M., Givens, B., & Bruno, J. F. (2001). The cognitive neuroscience of sustained attention: where top-down meets bottom-up. Brain Research Reviews, 35(2), 146–160. https://doi.org/10.1016/s0165-0173(01)00044-3

Schaefer, A., Kong, R., Gordon, E. M., Laumann, T. O., Zuo, X., Holmes, A. J., Eickhoff, S. B., & Yeo, B. T. (2018b). Local-Global Parcellation of the Human Cerebral Cortex from Intrinsic Functional Connectivity MRI. Cerebral Cortex, 28(9), 3095–3114. https://doi.org/10.1093/cercor/bhx179

Seli, P., Cheyne, J. A., & Smilek, D. (2013). Wandering minds and wavering rhythms: Linking mind wandering and behavioral variability. Journal of Experimental Psychology: Human Perception and Performance, 39(1), 1–5. https://doi.org/10.1037/a0030954

Siegel, M., Donner, T. H., & Engel, A. (2012). Spectral fingerprints of large-scale neuronal interactions. Nature Reviews Neuroscience, 13(2), 121–134. https://doi.org/10.1038/nrn3137

Smith, S. M., Jenkinson, M., Woolrich, M. W., Beckmann, C. F., Tej, B., Johansen-Berg, H., Bannister, P. R., De Luca, M., Drobnjak, I., Flitney, D., Niazy, R. K., Saunders, J., Vickers, J. C., Zhang, Y., De Stefano, N., Brady, J. N., & Matthews, P. M. (2004). Advances in functional and structural MR image analysis and implementation as FSL. NeuroImage, 23, S208– S219. https://doi.org/10.1016/j.neuroimage.2004.07.051

Spreng, R. N., & Grady, C. L. (2010). Patterns of Brain Activity Supporting Autobiographical Memory, Prospection, and Theory of Mind, and Their Relationship to the Default Mode Network. Journal of Cognitive Neuroscience, 22(6), 1112–1123. https://doi.org/10.1162/jocn.2009.21282

Spreng, R. N., Mar, R. A., & Kim, A. Y. (2009). The Common Neural Basis of Autobiographical Memory, Prospection, Navigation, Theory of Mind, and the Default Mode: A Quantitative Meta-analysis. Journal of Cognitive Neuroscience, 21(3), 489–510. https://doi.org/10.1162/jocn.2008.21029

Spreng, R. N., Sepulcre, J., Turner, G. R., Stevens, W. D., & Schacter, D. L. (2013). Intrinsic Architecture Underlying the Relations among the Default, Dorsal Attention, and Frontoparietal Control Networks of the Human Brain. Journal of Cognitive Neuroscience, 25(1), 74–86. https://doi.org/10.1162/jocn_a_00281

Spyropoulos, G., Bosman, C. A., & Fries, P. (2018). A theta rhythm in macaque visual cortex and its attentional modulation. Proceedings of the National Academy of Sciences of the United States of America, 115(24). https://doi.org/10.1073/pnas.1719433115

Strakowski, S. M., Adler, C. M., Holland, S. K., Mills, N. P., & DelBello, M. P. (2004). A Preliminary fMRI Study of Sustained Attention in Euthymic, Unmedicated Bipolar Disorder. Neuropsychopharmacology, 29(9), 1734–1740. https://doi.org/10.1038/sj.npp.1300492

Tamm, L., Narad, M. E., Antonini, T. N., O’Brien, K. D., Hawk, L. W., & Epstein, J. N. (2012). Reaction Time Variability in ADHD: A Review. Neurotherapeutics, 9(3), 500–508. https://doi.org/10.1007/s13311-012-0138-5

Thompson, G. J., Magnuson, M. L., Merritt, M. D., Schwarb, H., Pan, W., McKinley, A., Tripp, L. D., Schumacher, E. H., & Keilholz, S. D. (2013). Short-time windows of correlation between large-scale functional brain networks predict vigilance intraindividually and interindividually. Human Brain Mapping, 34(12), 3280–3298. https://doi.org/10.1002/hbm.22140

Thompson, G. J., Pan, W., Magnuson, M. L., Jaeger, D., & Keilholz, S. D. (2014). Quasi-periodic patterns (QPP): Large-scale dynamics in resting state fMRI that correlate with local infraslow electrical activity. NeuroImage, 84, 1018–1031. https://doi.org/10.1016/j.neuroimage.2013.09.029

Unsworth, N., & Robison, M. K. (2017). A locus coeruleus-norepinephrine account of individual differences in working memory capacity and attention control. Psychonomic Bulletin & Review, 24(4), 1282–1311. https://doi.org/10.3758/s13423-016-1220-5

Valera, F., Toro, A., John, E. R., & Schwartz, E. L. (1981). Perceptual framing and cortical alpha rhythm. Neuropsychologia, 19(5), 675–686. https://doi.org/10.1016/0028-3932(81)90005-1

VanRullen, R., & Koch, C. (2003). Is perception discrete or continuous? Trends in Cognitive Sciences, 7(5), 207–213. https://doi.org/10.1016/s1364-6613(03)00095-0

Vincent, J. L., Kahn, I., Snyder, A. Z., Raichle, M. E., & Buckner, R. L. (2008). Evidence for a Frontoparietal Control System Revealed by Intrinsic Functional Connectivity. Journal of Neurophysiology, 100(6), 3328–3342. https://doi.org/10.1152/jn.90355.2008

Von Stein, A., & Sarnthein, J. (2000). Different frequencies for different scales of cortical integration: from local gamma to long range alpha/theta synchronization. International Journal of Psychophysiology, 38(3), 301–313. https://doi.org/10.1016/s0167-8760(00)00172-0

Vossel, S., Geng, J. J., & Fink, G. R. (2014). Dorsal and Ventral Attention Systems. The Neuroscientist, 20(2), 150–159. https://doi.org/10.1177/1073858413494269

Weissman, D. H., Roberts, K. C., Visscher, K. M., & Woldorff, M. G. (2006). The neural bases of momentary lapses in attention. Nature Neuroscience, 9(7), 971–978. https://doi.org/10.1038/nn1727

Woolrich, M. W., Jbabdi, S., Patenaude, B., Chappell, M. A., Makni, S., Behrens, T. E., Beckmann, C. F., Jenkinson, M., & Smith, S. M. (2009). Bayesian analysis of neuroimaging data in FSL. NeuroImage, 45(1), S173–S186. https://doi.org/10.1016/j.neuroimage.2008.10.055

Xu, N., Smith, D. M., Jeno, G., Seeburger, D. T., Schumacher, E. H., & Keilholz, S. D. (in press). The interaction between random and systematic visual stimulation and infraslow quasiperiodic spatiotemporal patterns of whole brain activity. Imaging Neuroscience. bioRxiv link: https://www.biorxiv.org/content/10.1101/2022.12.06.519337v4

Yeo, B. T., Krienen, F. M., Sepulcre, J., Sabuncu, M. R., Lashkari, D., Hollinshead, M. O., Roffman, J. L., Smoller, J. W., Zöllei, L., Polimeni, J. R., Fischl, B., Liu, H., & Buckner, R. L. (2011). The organization of the human cerebral cortex estimated by intrinsic functional connectivity. Journal of Neurophysiology, 106(3), 1125–1165. https://doi.org/10.1152/jn.00338.2011

Yousefi, B., & Keilholz, S. D. (2021). Propagating patterns of intrinsic activity along macroscale gradients coordinate functional connections across the whole brain. NeuroImage, 231, 117827. https://doi.org/10.1016/j.neuroimage.2021.117827

Yousefi, B., Shin, J., Schumacher, E. H., & Keilholz, S. D. (2018). Quasi-periodic patterns of intrinsic brain activity in individuals and their relationship to global signal. NeuroImage, 167, 297–308. https://doi.org/10.1016/j.neuroimage.2017.11.043

Zuberer, A., Kucyi, A., Yamashita, A., Wu, C. M., Walter, M., Valera, E. M., & Esterman, M. (2021). Integration and segregation across large-scale intrinsic brain networks as a marker of sustained attention and task-unrelated thought. NeuroImage, 229, 117610. https://doi.org/10.1016/j.neuroimage.2020.117610

